# Structural and biochemical insights reveal substrate-modulated nuclease activity of ComEC during DNA processing

**DOI:** 10.64898/2026.04.23.720307

**Authors:** Sophie Deselaers, Dianhong Wang, Tamino Cairoli, Pavel Afanasyev, Manuela K. Hospenthal

## Abstract

Natural transformation enables bacteria to internalise extracellular DNA, driving adaptation and the spread of antibiotic resistance. The membrane protein ComEC mediates translocation of single-stranded DNA across the cytoplasmic membrane while degrading the complementary strand, yet the structural basis of its activity remains incompletely defined. Here, we report a cryo-electron microscopy structure of full-length ComEC from *Neomoorella carbonis* in a pre-translocation state, revealing a three-domain architecture and a conserved transmembrane channel captured in a closed conformation. Structural analysis indicates that conformational rearrangements of channel-lining helices are required to accommodate single-stranded DNA. Biochemical assays show that, relative to the isolated β-lactamase-like domain, full-length ComEC degrades DNA more efficiently. Importantly, coating of the DNA by the periplasmic DNA receptor ComEA suppresses endonucleolytic cleavage, thereby modulating nuclease activity. Together, these findings provide the first characterisation of the nuclease activity of full-length ComEC and show how ComEA-mediated protection of the substrate directs ComEC’s nuclease activity to ensure high fidelity during the natural transformation process.

## Introduction

Many bacterial species are able to internalise extracellular DNA (transforming DNA; tDNA) from their environment through natural transformation, a horizontal gene transfer pathway that shapes evolutionary adaptation and contributes to the dissemination of antibiotic resistance (Arnold *et al*, 2022; Dubnau & Blokesch, 2019; Winter *et al*, 2021). This process is mediated by a coordinated set of proteins that mediate DNA capture, processing and translocation, and integration.

Type IV pili or related structures mediate initial DNA binding (Piepenbrink, 2019), before transporting it into the periplasmic space by a mechanism that remains poorly understood. In Gram-positive organisms, the DNA is subsequently bound by the membrane-anchored receptor ComEA, which is composed of an N-terminal transmembrane helix, a central oligomerisation domain and a C-terminal atypical helix-hairpin-helix DNA binding domain (Ahmed *et al*, 2022). ComEA from *Geobacillus stearothermophilus* was shown to exist in a monomer-dimer equilibrium in solution (Ahmed *et al*, 2022), while in the presence of DNA, ComEA forms oligomers along its DNA substrate through the oligomerisation domain (Santiago *et al*, 2026).

DNA translocation across the plasma membrane is mediated by the channel protein ComEC, which is indispensable for natural transformation (Facius, 1993; Inamine & Dubnau, 1995; Pestova & Morrison, 1998). ComEC translocates a single strand of DNA across the membrane (Inamine & Dubnau, 1995), while the second strand is degraded by nuclease activity. In many competent organisms, this enzymatic activity is harboured within the β-lactamase-like domain encoded at the C-terminus of ComEC (Baker *et al*, 2016; Silale *et al*, 2021). In addition to the β-lactamase-like domain, most ComEC orthologues contain an oligonucleotide binding (OB) domain encoded near the N-terminus and a central channel/competence domain (Baker *et al*, 2016; Pimentel & Zhang, 2018). Recently we determined the structure of the isolated β-lactamase-like domain from *Moorella glycerini* (recently proposed as *Neomoorella glycerini* (Gtari & Ventura, 2025)) and showed that it functions as an endo- and 5’ exonuclease *in vitro* (Stedman *et al*, 2025). Our structural and biochemical observations led us to propose a model of 5’ strand-specific topological processivity of the β-lactamase-like domain, which in turn prevents the degradation of the opposite 3’ strand, leading to the translocation of the latter across the membrane. This is consistent with previous work that determined the polarity of DNA transport across the plasma membrane (Méjean & Claverys, 1988). Once the ssDNA emerges in the cytoplasm, the ATP-dependent DNA translocase ComFA contributes by potentially exerting a pulling force on the DNA (Foster *et al*, 2022).

However, whether endonucleolytic cleavage events by the β-lactamase-like domain occur *in vivo*, and if so, what significance this has, remains unclear. Furthermore, the activity of the nuclease domain in the context of full-length ComEC also remains untested. The lack of biochemical, biophysical, and structural characterization of ComEC has been a major limitation in understanding how ComEA transfers DNA to ComEC, and how ComEC subsequently binds, cleaves, and translocates DNA. Any such efforts have in turn been hindered by the lack of access to the full-length ComEC protein for *in vitro* studies.

To address this, we determined a cryogenic electron microscopy (cryo-EM) structure of full-length ComEC from *Moorella carbonis* (recently proposed as *Neomoorella carbonis* (Gtari & Ventura, 2025)). Our structure provides a complete view of ComEC, defining the organisation of the transmembrane helices that form the conserved DNA channel in the competence domain. Through modelling of the translocation complex, we propose the conformational rearrangements required to enable ssDNA translocation through the channel. In addition to our structural analysis, we compared the nuclease activity of full-length ComEC with that of the isolated β-lactamase-like domain and tested whether its cleavage mode is affected by the coating of DNA with ComEA. We find that full-length ComEC has markedly enhanced nuclease activity and that ComEA modulates access of the nuclease domain to its DNA substrate, thereby influencing whether cleavage proceeds via internal cuts. Together, these results establish a structural framework for full-length ComEC and reveal how ComEA shapes the enzymatic processing of tDNA, providing mechanistic insight into how DNA processing is regulated during natural transformation.

## Results

### The cryo-EM structure of *apo* ComEC

We set out to identify a ComEC orthologue that expresses and purifies sufficiently well for biochemical and structural analyses. We screened >20 ComEC orthologues, including thermophilic organisms and close relatives of *N. glycerini*, which has previously enabled our structural work of ComEC’s β-lactamase-like and OB domains (Stedman *et al*, 2025). We recombinantly expressed full-length *N. carbonis* ComEC (ComEC_Nc_) in *Escherichia coli*, purified the sample by affinity and size exclusion chromatography (**Fig. S1**), and applied it to grids which were vitrified for cryo-EM analysis (**Fig. S2**). We obtained two maps, of full-length ComEC and the locally refined β-lactamase-like domain, resolved at 4.1 and 4.2 Å, respectively. This allowed a composite model of ComEC in its entirety to be unambiguously built and refined (**Fig. 1a, b, Fig. S2, Table S1**). In addition to the periplasmically located OB fold and the C-terminal β-lactamase-like domain, the ComEC_Nc_ competence domain contains 11 transmembrane (TM) helices, three of which occur before the OB fold, as well as three long and four short helices that do not span the membrane (non-spanning (NS) helix) (**Fig. 1b, c, Fig. S3**). The first non-membrane spanning helix, lying approximately parallel to the membrane plane (NS1), is an amphipathic helix that immediately follows the OB fold. This is followed by two shorter and slightly more tilted periplasmic helices (NS2 and NS3). Additionally, short helices are located between NS3 and TM4 (NS4) and between TM7 and TM8 (NS5), with NS4 embedded within the membrane and NS5 positioned at the membrane interface. Between TM9 and TM10, two additional membrane-parallel and non-spanning helices (NS6 and NS7) are largely buried within the lipid bilayer. Two short β-strands immediately before and after NS6 and NS7, respectively, come together forming a small anti-parallel β-sheet. NS1-4, NS6, and NS7 arrange into a peripheral helical bundle forming a distinct structural unit on one side of the protein. Lastly, there is a short cytoplasmic helix (CH1) located between TM10 and TM11. Within the core transmembrane region, a putative ssDNA channel can be seen forming a bent, yet continuous narrow pore through the competence domain. To our surprise, ComEC formed an asymmetric dimer in our structure (**Fig. S4**), which may reflect a sample preparation-induced artefact. The two ComEC protomers in the dimer adopt a highly similar overall structure, except for a subtle rigid body movement of the entire β-lactamase-like domain and minor local differences (**Movie S1**). This breathing motion of the β-lactamase-like domain limits the resolution of the map in this region compared to the OB fold and the competence domain. In addition, the quality of the map differs between the two ComECs of the dimer, with TM11 entirely absent in one protomer. Each ComEC protomer appears functionally complete and likely capable of DNA capture, processing and translocation.

**Fig. 1:**
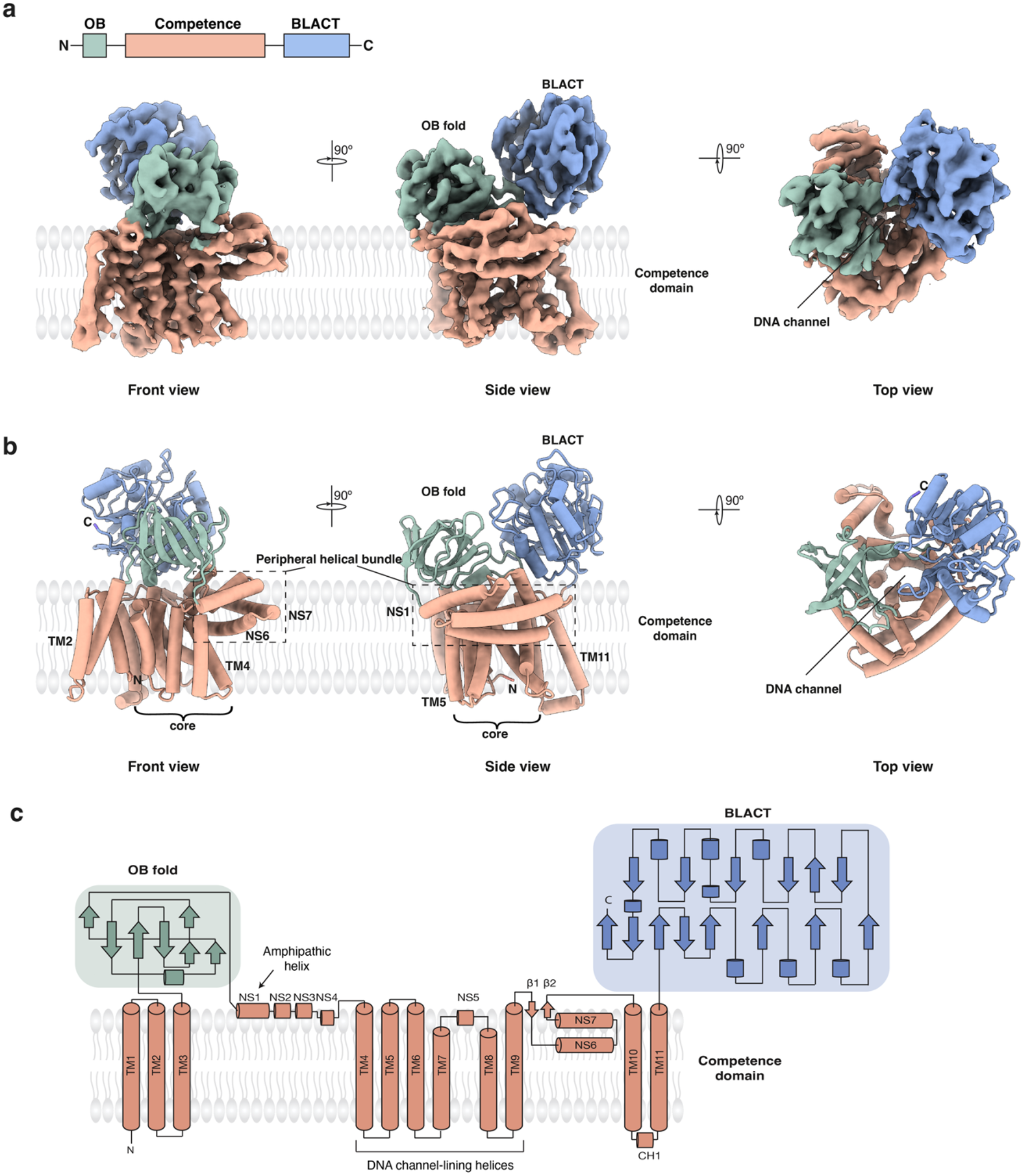
The cryo-EM structure of ComEC. **a** Top: diagram illustrating the domain organisation of ComEC_Nc_. Bottom: Three views of the cryo-EM map of ComEC_Nc_ coloured by domain, with the OB fold shown in green, the β-lactamase-like domain in blue and the transmembrane competence domain in orange. **b** The atomic model of ComEC_Nc_ shown in the same three views. The DNA channel and the peripheral helical bundle are indicated on the figure. **c** The corresponding topology diagram illustrating all secondary structural elements and domains. The amphipathic helix and the channel-lining helices are also labelled.

### tDNA translocation requires conformational changes in the competence domain

To understand tDNA translocation by ComEC, we analysed the DNA channel in the competence domain in further detail. The DNA pathway through ComEC can be subdivided into the periplasmic entrance, between the OB fold and β-lactamase-like domain, and the membrane-spanning portion of the channel. In the *apo* state, the putative ssDNA channel is is surrounded by TM4-9, with most predicted channel-lining residues contributed by TM4, TM6, TM8, and TM9. The N-terminal part of NS6 also contributes to the channel, whereas TM1-3, as well as TM10 and TM11 are located more peripherally. First, we examined the level of conservation of the predicted channel-lining residues (**Fig. 2a, Fig S5**). The majority of channel-lining residues within the membrane-spanning region of the channel are highly conserved, whereas some residues at the periplasmic entrance (R99, Y101 and F572) and channel exit (Y319, F327 and D369) are less conserved. Next, we compared the channel in our *apo* cryo-EM structure with the AlphaFold3-predicted translocation complex containing ssDNA traversing the channel (Abramson *et al*, 2024). Comparison of the two models shows that the channel of the translocation complex adopts a straighter trajectory, whereas the channel in our experimentally determined *apo* structure follows a more curved path. This likely reflects a non-translocating, closed conformation in the *apo* structure. Next, we used the MOLEonline tool (Raček *et al*, 2025) to visualise the DNA channel in both our *apo* ComEC structure, as well as the predicted translocation complex (**Fig. 2a, b**). In the *apo* structure, the narrowest point of the channel is located just below the periplasmic membrane boundary and widens toward both the periplasmic entrance and the channel exit region. The smallest distance occurs between V258 and G350 (Cα distance 6.2 Å) (**Fig. 2c**). Given that an extended single DNA strand has an effective diameter of ~10 Å, this constriction would preclude ssDNA passage without conformational rearrangement, even without accounting for side chain contributions. In the predicted translocation complex, the channel is wider but retains a constriction of 10.2 Å (at the same position, Cα measurement), suggesting that yet further widening may be required for ssDNA translocation. Our comparative analysis suggests that channel widening is achieved by coordinated rearrangments of the same transmembrane helices that formed the predicted channel in the *apo* structure. This likely occurs via a displacement of NS4 and the adjoining NS4-TM4 loop (**Fig. 2c**), accompanied by lateral movements of TM4-6 (**Fig. 2c, d**). The predicted magnitude of these movements is largest in the channel exit region. In addition to the channel flanking helices, TM11 also undergoes a considerable conformational change (**Movie S2**). Taken together, our structural characterisation of ComEC reveals a DNA channel lined by highly conserved residues and defines the structural rearrangements required to accommodate and translocate tDNA.

**Fig. 2:**
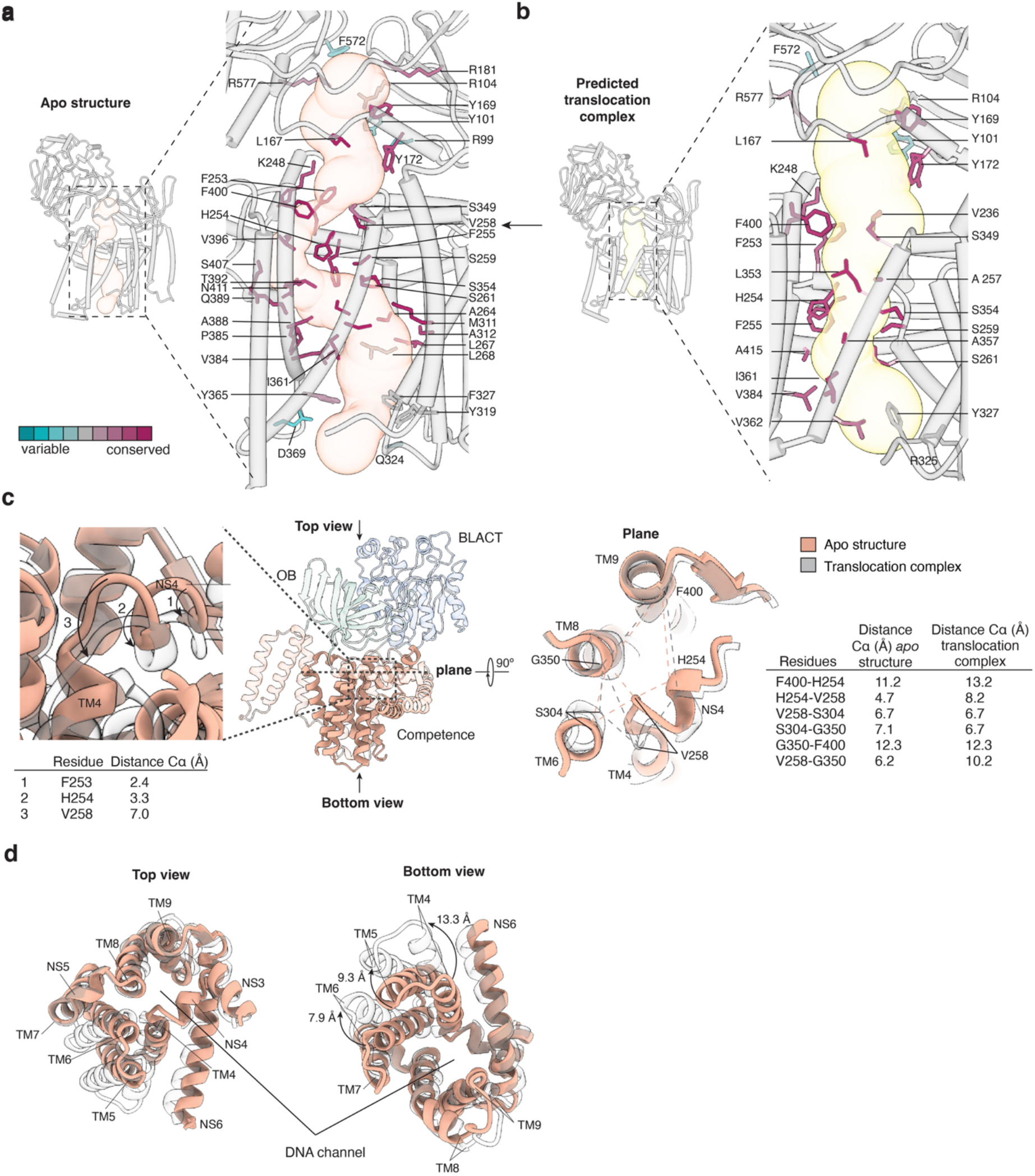
The channel undergoes conformational changes to translocate ssDNA. **a, b** Channel diagrams of the *apo* ComEC cryo-EM structure (**a**) and the AlphaFold3 prediction of the translocation complex (ssDNA was removed for visualisation purposes) (**b**) generated using MOLEonline. Channel-lining residues are coloured according to their conservation. For illustrative purposes, all channel-lining residues are shown, including those with insufficient density to define side chain orientations. An arrow indicates the narrowest point of the channel in the *apo* structure. **c** Comparison of the channel between our *apo* ComEC structure and the AlphaFold3 prediction the translocation complex. Left, close-up view of the boxed region in the orientation diagram showing the rearrangement of NS4 and the NS4-TM4 loop. Arrows show the movement of the Cα atoms of F253, H254 and V258 between the two models and distances are indicated. Right, the rearrangement of helices at the narrowest point of the channel between the two models. Distances between anchor Cα positions between structurally neighbouring helices are shown for both models. **d** Top and bottom views of the channel-lining helices and their movements between our *apo* structure and the AlphaFold model of the translocation complex.

### OB fold stabilisation in the full-length structure positions the active site of the nuclease domain at the channel entrance

Having defined the architecture of the transmembrane channel, we next investigated how the periplasmic domains are arranged in the full-length structure and how their interaction positions the active site with respect to the channel entrance. First, we compared our structural models of the β-lactamase-like domain and the OB fold within full-length ComEC with our previously determined structures of each isolated domain (Stedman *et al*, 2025) (**Fig. 3a, b**). The OB folds and β-lactamase-like domains from *N. glycerini* and *N. carbonis* share 90.3% and 92.5% sequence identity, respectively, and are therefore expected to adopt highly similar structures. Consistent with this, the β-lactamase-like domain remains largely unchanged, with limited flexibility in loop regions and a Cα root mean square deviation (RMSD) of 1.01 Å (over 232 aligned atoms) (**Fig. 3a**). In contrast, a small subset of loops in the OB fold that were previously unstructured adopt defined secondary structure in the full-length ComEC structure (**Fig. 3b**). Most notably, the loop between βIV and βV forms a short helical segment (αI). In the full-length structure, αI contacts NS5 of the competence domain and contains the highly conserved aromatic residues (Y169 and Y172), likely involved in stabilising the DNA substrate at the periplasmic entrance of the channel (**Fig. 3c**). In addition, the loop preceding αI of the OB fold extends to contact the β-lactamase-like domain, forming a small but conserved interface (**Fig. 3d**). We propose that this interface positions and orients the β-lactamase-like domain, which would otherwise remain mobile due to its flexible linker connection to the competence domain. In addition, residues from both domains contribute to the periplasmic entrance region of the DNA channel (**Fig. 2a**). Together, these observations indicate that structural ordering of the OB fold within the full-length context of ComEC establishes interdomain interactions that position the active site of the β-lactamase-like domain at the periplasmic channel entrance and contribute to substrate engagement.

**Fig. 3:**
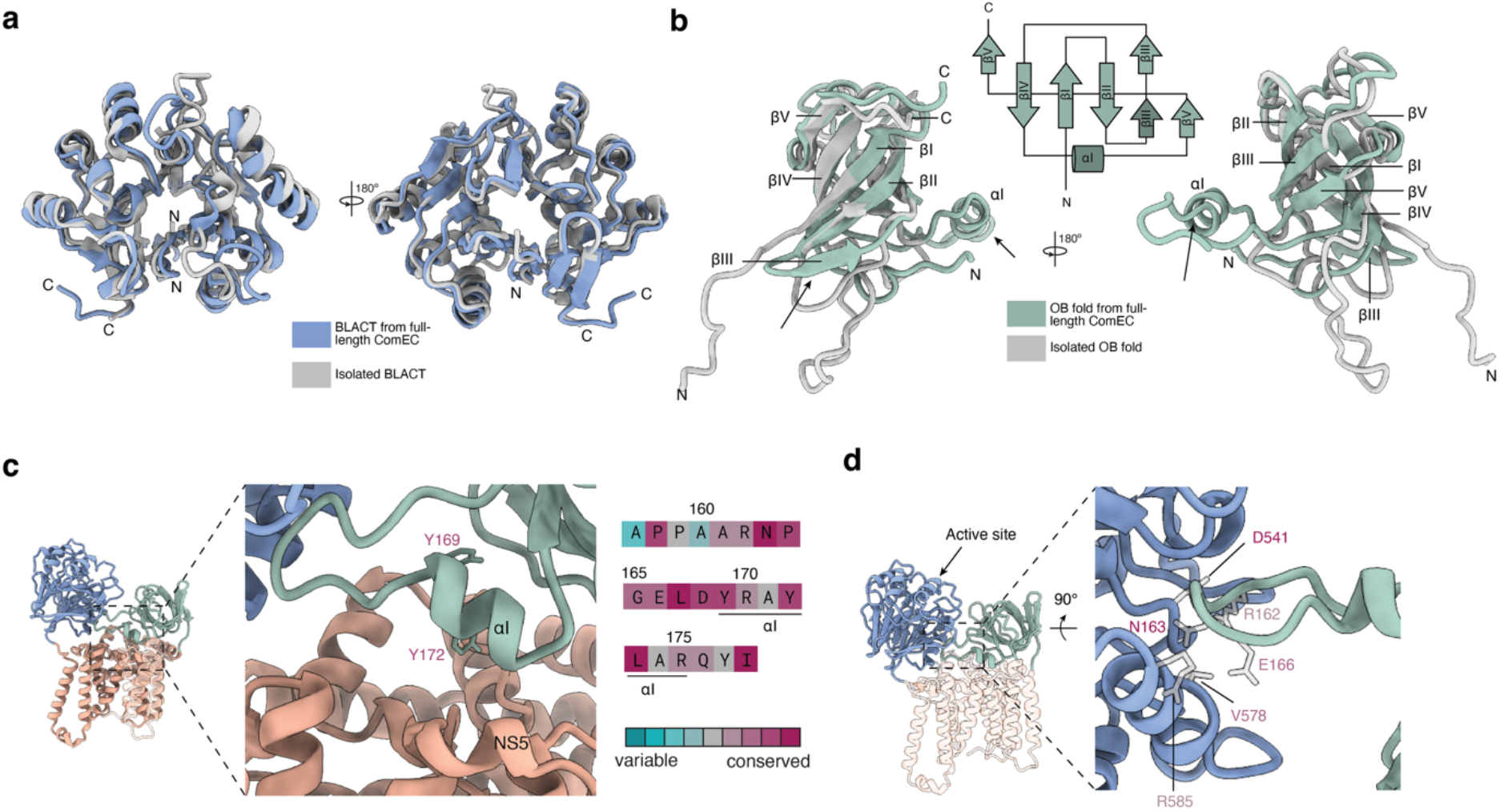
The interdomain interfaces shape ComEC’s architecture. **a** Superposition of the β-lactamase-like domain from our cryo-EM structure of full-length ComEC_Nc_ (blue) with the previously determined crystal structure of the isolated β-lactamase-like domain from *N. glycerini* (PDB ID: 9IC4) (grey). **b** Superposition of the OB fold from our cryo-EM structure of full-length ComEC_Nc_ (green) with the previously determined NMR structure of the isolated OB fold from *N. glycerini* (PDB ID: 9IEW) (grey) shown in two orientations. A topology diagram highlighting structural changes (darker shade) of the OB fold that occur in the context of full-length ComEC. **c** Close-up view of the αI helix in the OB fold that becomes ordered in the full-length context of ComEC. Aromatic residues at the periplasmic channel entrance are shown and their figure labels are coloured according to sequence conservation. The sequence of the βIV-βV loop region, including αI, coloured accoring to conservation. **d** Close-up view of the interface between the βIV-βV loop and the isolated β-lactamase-like domain. For illustrative purposes side chains are shown in grey, despite limiting density in this area, and their labels are coloured according to conservation. The position of the active site in the β-lactamase-like domain is marked by an arrow.

### Full-length ComEC architecture and ComEA modulate ComEC nuclease activity

Having defined the structural organisation of ComEC, we next examined whether nuclease activity differs between the isolated β-lactamase-like domain and the full-length protein, and how it is further modulated by ComEA-coated DNA. In our previous study, we showed that the β-lactamase-like domain from *N. glycerini* (BLACT_Ng_) displays both endo- and 5′ exonuclease activity, with activity enhanced upon association with the OB fold, and proposed a model of strand-specific topological processivity to explain how ComEC selectively degrades one strand while preserving the other (Stedman *et al*, 2025).

We reasoned that in the context of full-length ComEC, where all three domains adopt their native arrangement, nuclease activity may be further enhanced. Therefore, we performed time-course nuclease activity assays with FAM-labelled ssDNA at different temperatures and found maximal activity at ~60°C (**Fig. 4a**), consistent with the optimal growth temperature of *Neomoorella* species (Böer *et al*, 2024). All subsequent assays were therefore performed at this temperature. Next, we compared nuclease activity of full-length ComEC_Nc_ and the isolated β-lactamase-like domain (BLACT_Nc_) using ssDNA and dsDNA substrates (**Fig. 4b**). Full-length ComEC_Nc_ degrades ssDNA more rapidly than dsDNA and is overall more active than BLACT_Nc_. The more pronounced difference in degradation rates between ssDNA and dsDNA may reflect additional constraints imposed by duplex DNA and/or differences in cleavage mode.

**Fig. 4:**
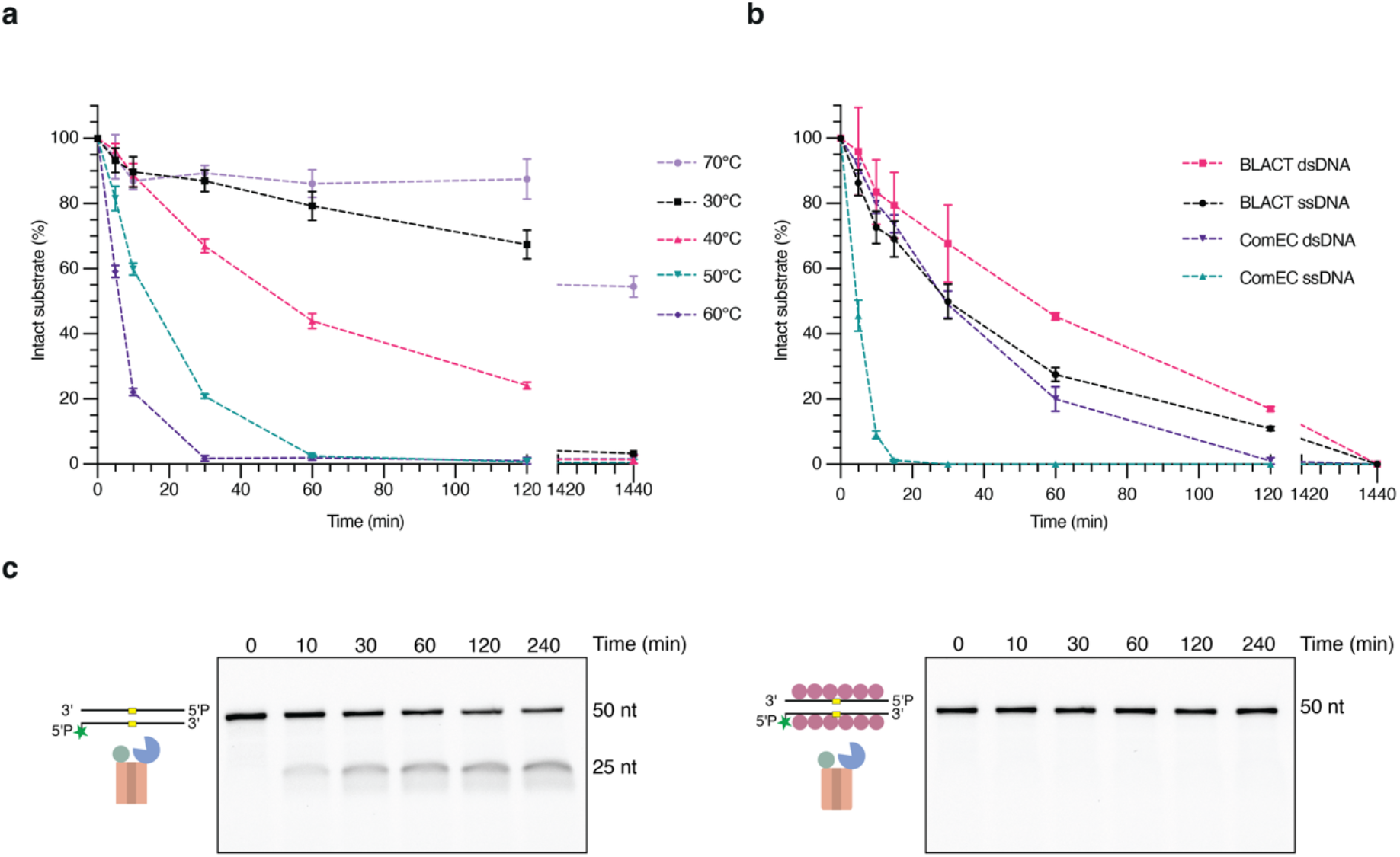
Full-length ComEC exhibits enhanced nuclease activity that is modulated by ComEA. **a**-**c** Nuclease activity assays monitoring the degradation of FAM-labelled DNA substrates by ComEC constructs. Error bars represent the standard error from three technical replicates. **a** Temperature dependence of ssDNA degradation by ComEC_Nc_. **b** Comparison of ssDNA and dsDNA degradation by ComEC_Nc_ and BLACT_Nc_. **c** Comparison of cleavage of naked and ComEA-coated dsDNA substrates containing defined nuclease-resistant PTO linkages and a single central cleavable phosphodiester bond (yellow rectangle). The 5′ ends of both strands are phosphorylated, and one strand carries a FAM label (green star).

We next tested whether coating of the DNA substrate with ComEA_Ng_, reflecting the physiological substrate, suppresses endonucleolytic cleavage by ComEC (**Fig. 4c, Fig. S6**). We used substrates containing a single internal cleavable phosphodiester bond flanked by nuclease-resistant phosphorothioate (PTO) linkages and a ComEA construct containing only the oligomerisation and DNA binding domains. Using pre-formed ComEA-DNA filaments as the substrate, we observed a near-complete suppression of endonucleolytic products compared to naked DNA. This suggests that ComEA limits access of the nuclease active site to the DNA backbone. Together, these results show that nuclease activity is intrinsically enhanced in the full-length context, and is further constrained by ComEA-coated DNA to suppress endonucleolytic cleavage. ComEA may contribute to transformation fidelity by limiting unrestrained endonucleolytic cleavage that may otherwise target the translocating strand.

## Discussion

ComEC is essential for natural transformation in both Gram-positive and Gram-negative bacteria, yet the molecular basis of DNA translocation across the cytoplasmic membrane has remained poorly defined. Here, we present a cryo-EM structure of full-length *apo* ComEC from *Neomoorella carbonis*, combined with structural modelling of conformational changes required for ssDNA passage. Furthermore, we characterise ComEC’s nuclease activity and uncover a novel role for ComEA in DNA processing, extending its function beyond DNA uptake.

The ComEC structure reveals a three-domain architecture, with the OB fold and β-lactamase-like domain positioned adjacent to one another on the periplasmic side of the membrane (**Fig. 1**). Within the competence domain, TM4-9 form a highly conserved ssDNA channel that, even in the *apo* state revealed by our structure, forms a continuous but constricted pathway across the membrane. Structural analysis of the channel suggests that substantial conformational rearrangements are required to support DNA translocation (**Fig. 2**). Comparison of our *apo* structure with an AlphaFold predition of a complex translocating DNA, shows that the channel straightens and widens during this process. At the periplasmic entrance, the NS4-TM4 loop is predicted to undergo a substantial rearrangement, moving downward and outward to relieve partial occlusion of the channel. In addition, our modelling shows lateral displacement of TM4-6, particularly near the cytoplasmic channel exit, resulting in widening of the channel. Despite these conformational changes, the predicted translocation complex still contains constrictions that appear too narrow to accommodate ssDNA, indicating that the AlphaFold model of the translocation complex likely underestimates the required conformational rearrangments. Another interesting feature of our structure is that ComEC forms an asymmetric dimer, in which local conformational differences are observed between protomers and the β-lactamase-like domain exhibits mobility (**Fig. S4**). However, ComEC likely functions as a monomer *in vivo*, as a single protomer appears sufficient to permit DNA binding, degradation and translocation.

Membrane protein channels often employ gating mechanisms to restrict the unwanted passage of small molecules and ions (Drew & Boudker, 2015). We therefore analysed our *apo* structure and the modelled translocation complex to identify which regions of ComEC may contribute to potential gating (**Fig. 2c, Fig. S7**). In the *apo* structure, the NS4-TM4 loop partially occludes the channel and the adjacent NS4 helix contains several conserved bulky aromatic residues (F253, H254, F255 and F256). This loop undergoes predicted conformational changes upon DNA translocation. Inspection of the side-chain density in this region across both protomers of the ComEC dimer revealed conformational variability for H254, which adopts at least two distinct orientations. In one conformation, the channel is more constricted in the vicinity of H254. These observations are consistent with a potential role for this region in gating. In contrast, the cytoplasmic exit region of the channel appears comparatively wide and lacks an obvious structural element capable of occlusion. After engagement of the DNA substrate, such a gate would need to open to permit translocation. In addition, ComEC is unlikely to be constitutively expressed and may be expressed mostly during the competence state, providing an additional level of control to limit unintended permeability.

Comparison of the full-length ComEC structure with previously determined structures of the isolated OB fold and β-lactamase-like domain reveals important conformational changes within the OB fold (**Fig. 3**). The βIV-βV loop, which is fully disordered in our previous NMR structure (Stedman *et al*, 2025), now includes a short α-helical segment (αI) and contributes to a conserved interface with the β-lactamase-like domain (**Fig. 3d**). Although the precise contacts are not fully resolved, the presence of conserved charged residues suggest salt bridge interactions, while the limited extent of the interface indicates that it is relatively weak. Such an interaction might be sufficiently strong to orient the nuclease active site appropriately for DNA engagement at the periplasmic channel entrance region, while retaining the conformational flexibility that may be required for DNA accommodation and translocation.

Biochemical analysis reveals that full-length ComEC degrades DNA more efficiently than the isolated β-lactamase-like domain (**Fig. 4b**), indicating that the native domain organisation enhances nuclease activity. Furthermore, we observed a more pronounced difference in degradation rates between ssDNA and dsDNA by full-length ComEC. This may be due to additional constraints imposed by threading the non-degraded strand through the DNA channel during translocation.

A key finding of this study is that ComEA, previously implicated to function predominantly in the DNA uptake step, also affects DNA processing by ComEC. We show that coating of DNA with ComEA, mimicking the physiological substrate of ComEC, strongly suppresses ComEC’s endonucleolytic cleavage (**Fig. 4c**). By limiting access of the nuclease active site to the DNA backbone, ComEA is thus able to modulate ComEC’s cleavage mode. This may be functionally important *in vivo*, as unrestrained endonucleolytic activity may otherwise cleave the translocating strand. In this context, ComEA may play a critical role in safeguarding the fidelity of natural transformation. Endonucleolytic suppression by ComEA is likely most effective within internal regions of the DNA substrate, whereas cleavage near DNA ends may still occur due to less complete ComEA coating and transient exposure of the DNA backbone. Such a mechanism would be consistent with previous studies reporting mono-, di- and trinucleotide products *in vivo* (Claverys *et al*, 2009). This is also in line with our previously proposed model of strand-specific topological processivity, in which ComEC degrades the 5′ leading strand while threading the 3′ strand through the DNA channel (Stedman *et al*., 2025). While remaining engaged with the substrate, ComEC may employ exonucleolytic, endonucleolytic (producing short fragments), or mixed cleavage modes to processively degrade the 5′ strand. The relative contributions of these activities, as well as their coordination with DNA translocation remain to be determined.

Important questions regarding the mechanism of ComEC remain. Recent structural work has reported ComEC structures with DNA bound at the periplasmic entrance (Hirano *et al*, 2026). However, a complete mechanistic understanding will require additional structural information capturing ComEC engaged in DNA degradation and DNA translocation.

In summary, we report a cryo-EM structure of ComEC in a pre-translocation state, model the structural rearrangments required for translocation, and show that nuclease activity is enhanced in the full-length protein and modulated by ComEA to enable controlled DNA processing during natural transformation.

## Methods

### Plasmids

The ComEC and BLACT constructs are based on the sequences encoded in *Neomoorella carbonis*, while the ComEA construct is derived from the closely related *Neomoorella glycerini* (sequence identity = 79.2%). Full-length ComEC was recombinantly expressed from a pBAD vector harbouring an N-terminal sfGFP tag and a C-terminal StrepII tag and a 3C cleavage site following the sfGFP tag, whereas the β-lactamase-like domain (residues 532 to 801) was encoded on a pOPINS vector, with an N-terminal His_6_-Sumo tag. The ComEA construct consisted of the DNA binding and oligomerisation domain, and was expressed from a pET28 backbone encoding an N-terminal His_6_-Sumo tag. In-Fusion cloning was carried out according to the manufacturer’s guidelines with the HiFi PCR premix (Takara). Plasmids and Primers used in this study can be found in **Tables S2** and **S3**. DNA sequences were synthesised by Twist Bioscience.

### Protein production

ComEC_Nc_ was expressed in *Escherichia coli* C43 cells (New England Biolabs, NEB) grown in Luria–Bertani (LB) medium at 37°C and induced at an optical density at 600 nm (OD_600_) of~0.8 with 0.01% (w/v) arabinose for 4 h. Cells were resuspended in buffer A (50 mM HEPES-NaOH pH 7.5, 150 mM NaCl, 2 mM β-mercaptoethanol) supplemented with lysozyme (0.2 mg/mL), DNase I (10 µg/mL), and one cOmplete mini ethylenediaminetetraacetic acid (EDTA)-free protease inhibitor tablet (Roche), and lysed using an EmulsiFlex-C5 homogenizer (Avestin). Lysates were clarified by centrifugation (20’000 × *g*, 60 min) and membranes isolated by ultracentrifugation (40’000 rpm, 120 min, Ti45 rotor) (Beckman Coulter). Membranes were solubilised overnight in buffer A containing 1.5% (w/v) n-dodecyl-β-D-maltoside (DDM) with simultaneous 3C protease cleavage of the sfGFP tag. After ultracentrifugation (40’000 rpm, 60 min), the protein was purified by StrepTrap XT (Cytiva) affinity chromatography in buffer A supplemented with 0.01% (w/v) lauryl maltose neopentyl glycol (LMNG). ComEC_Nc_ was eluted with 50 mM biotin, concentrated, and further purified by size exclusion chromatography (Superdex 200 Increase 10/300, Cytiva) in buffer A.

BLACT_Nc_ and ComEA_Ng_ were expressed in *Escherichia coli BL21 (DE3)* cells (NEB) grown in LB medium at 37°C and induced at an OD_600_ of ~0.8 with 0.5 mM isopropyl-β-D-thiogalactoside (IPTG), followed by incubation at 18°C for 16–18 h. Cells expressing BLACT_Nc_ were resuspended and lysed as above, except in buffer B (50 mM HEPES-NaOH pH 7.2, 1 M NaCl, 20 mM imidazole, 2 mM β-mercaptoethanol). The clarified lysate was applied to a HisTrap HP column (Cytiva) and eluted with a 20–500 mM imidazole gradient. The His_6_-SUMO tag was removed by SENP1 during dialysis against buffer C (50 mM HEPES-NaOH pH 7.2, 50 mM NaCl, 2 mM β-mercaptoethanol). The protein was further purified by ion exchange chromatography (HiTrap Q, Cytiva) and size exclusion chromatography (Superdex 75 Increase 10/300, Cytiva) in buffer C. To purify ComEA_Ng_, buffer D (50 mM HEPES-NaOH pH 7.2, 150 mM NaCl, 10 mM imidazole) was used for resuspension. ComEA_Ng_ was bound to TALON Superflow resin (Cytiva). The resin was washed with buffer E (50 mM HEPES-NaOH pH 7.2, 1 M NaCl), incubated with benzonase, and washed again in buffer E. The protein was eluted from the beads by addition of SENP1 and further purified by size exclusion chromatography. The absorption at 280 nm combined with the corresponding molar extinction coefficients were used to determine protein concentrations (128690 M^-1^ cm^-1^ for ComEC_Nc_, 28420 M^-1^ cm^-1^ for BLACT_Nc_, 2980 M^-1^ cm^-1^ for ComEA_Ng_).

### Cryo-EM

#### Sample preparation and data collection

Cryo-EM samples were prepared by applying 3.5 µl of purified ComEC_Nc_ at a concentration of 5 mg/mL onto glow-discharged Quantifoil R1.2/1.3 holey carbon/copper 300 mesh grids. The grids were plunge-frozen in a liquid ethane and propane mixture using a Vitrobot Mark IV (Thermo Fisher Scientific). Data were collected on a Titan Krios G3i transmission electron microscope operated at 300 kV (Thermo Fisher Scientific), equipped with a K3 direct electron detector (Gatan) operated in counting mode and a BioContinuum energy filter (Gatan) with a slit width of 20 eV. Movies were recorded using EPU software (version 3.10.0) at a calibrated pixel size of 0.51 Å over a target defocus range of −1.1 to −2.7 µm. A total exposure of 53 electrons/Å^2^ was distributed over 40 frames.

#### Image processing and reconstruction

The single particle analysis workflow is summarised in **Fig. S2**. A gain reference was generated *a posteriori* from 1’000 movie stacks using Relion 5.0 (Burt *et al*, 2024; Afanasyev *et al*, 2015). All raw movie stacks and the gain reference were then imported into cryoSPARC v4.7.1 (Punjani *et al*, 2017). The frames of the movie stacks were motion-corrected and dose-weighted using patch-based motion correction at a binning factor of 2, resulting in a pixel size of 1.02 Å. The contrast transfer function (CTF) was estimated using patch CTF. A total of 89’904 micrographs were selected for further processing based on CTF estimation quality, defocus values and other parameters. Initial particle picking was performed on a subset of 20’076 micrographs using Blob picker. Particles were extracted with a box size of 320 pixels, binned to a pixel size of 4.08 Å, and subjected to 2D classification. Particles belonging to well-defined 2D class averages were selected for Topaz training (Bepler *et al*, 2020). The trained Topaz model was then used to pick particles from the entire dataset. A total of 3’276’527 particles were extracted (box size 320 pixels) and binned to a pixel size of 4.08 Å for 2D classification. Manually seleted 2D class averages were used to generate initial models for heterogenous refinement. Representative classes corresponding to larger particles, smaller particles, as well as “bad” particles (false-positive picks), were manually selected and used for *ab initio* reconstruction. These 3D volumes were used as initial models for iterative heterogeneous refinement. After initial cleaning of the dataset by heterogeneous refinement, 2D classification, and non-uniform refinement, 1’037’602 particles corresponding to the ComEC dimer were re-extracted at a pixel size of 1.53 Å and subjected to further heterogeneous refinement using one well-resolved class and three decoy classes to remove residual poorly aligned particles. Non-uniform refinement indicated that ComEC adopts an asymmetric dimer conformation. To preserve potential asymmetry, subsequent reconstructions were performed without imposing symmetry (C1). A subset of 480’597 high-quality particles was re-extracted without binning (pixel size 1.02 Å) and subjected to 3D classification without alignment to isolate the most homogeneous population. Subsequent local refinement yielded a reconstruction of the ComEC dimer at an estimated resolution of~4.1 Å, without symmetry imposed (C1). As the local resolution map indicated lower resolution in the β-lactamase-like domain, a focused refinement strategy was employed to improve map quality. For each dimeric particle, two monomeric sub-particles were generated by recentering on individual monomers and aligning them relative to one another. To minimise contributions from other regions, densities corresponding to the transmembrane (TM) and OB fold domains were removed by signal subtraction. A soft mask was generated in UCSF ChimeraX (Pettersen *et al*, 2021) using the “Cube’n Tube” plugin (Cairoli, 2026) . Signal-subtracted particles were subjected to 3D classification without alignment to resolve structural heterogeneity, followed by local refinement of the β-lactamase-like domain, yielding a reconstruction at ~4.2 Å resolution.

#### Model building and refinement

The cryo-EM density for the TM and OB fold domains within the dimeric ComEC map, as well as the β-lactamase-like domain obtained by focused refinement, provided sufficient detail for model building (**Fig. S2**). An initial model of monomeric ComEC was generated using AlphaFold3 (Abramson *et al*, 2024). For the dimeric ComEC map, the β-lactamase like domain was truncated from the predicted model, and two copies of the TM and OB fold domains were rigid-body fitted into the density. The model was manually adjusted in Coot (Emsley *et al*, 2010) and refined against the sharpened map using real-space refinement in PHENIX (Adams *et al*, 2010) and RosettaCM (Song *et al*, 2013). Refinement parameters in PHENIX included simulated annealing, reference restraints, and secondary structure restraints. Ramachandran and rotamer outliers were corrected manually in Coot.

A similar approach was used to build the β-lactamase-like domain model into the 4.2 Å map obtained by focused refinement. To generate the composite ComEC dimer model, the refined β-lactamase like domain model was docked into the dimeric ComEC map and combined with the TM and OB fold domains. The final full-length ComEC dimer model was subjected to an additional round of real-space refinement. Model quality assessment and model vs. map Fourier Shell Correlations (FSCs) calculations were performed using MolProbity and phenix.mtriage in PHENIX, respectively (Afonine *et al*, 2018; Adams *et al*, 2010). Refinement and validation statistics are summarised in **Table S1**. Figures were prepared using UCSF ChimeraX (Pettersen *et al*, 2021).

Global resolutions of cryo-EM density maps were estimated using the FSC=0.143 criterion (Rosenthal & Henderson, 2003). Model vs. map FSC curves were calculated, and resolutions are reported at the FSC=0.5 cutoff (Chen *et al*, 2013). Model validation statistics were assessed using MolProbity.

#### Nuclease activity assays

Nuclease activity was assessed using various fluorescein (FAM)-labelled DNA substrates (Microsynth; **Table S3**). Linear FAM-labelled dsDNA was prepared by annealing a FAM-labelled strand with its unlabelled complementary strand. Reactions were carried out in 50 mM HEPES-NaOH pH 7.2, 50 mM NaCl, 5 mM MnCl_2_, and 2 mM β-mercaptoethanol at 60°C, unless stated otherwise. At the indicated time points, aliquots were removed and quenched by mixing 1:1 with Novex 2x Tris-borate-EDTA (TBE)-urea sample buffer (Invitrogen). DNA fragments were separated on 12% denaturing polyacrylamide gels containing 7 M urea and visualized by fluorescence illumination using a ChemiDoc imaging system (Bio-Rad). Band intensities corresponding to the intact substrate were quantified and normalised to t = 0 min, and the fraction of remaining substrate was plotted over time. For Fig. 4a and b, reactions contained 0.2 µM enzyme and 10 µM ssDNA or 5 µM dsDNA. For Fig. 4c, substrate concentrations were adjusted to 2.5 µM and the DNA substrate was modified with phosphorothioate (PTO) linkages to visualise endonucleolytic activity, with ComEA_Ng_ added at a final concentration of 62.5 µM.

#### Electrophoretic mobility shift assay

To determine the concentration of ComEA required to fully saturate dsDNA fragments, an electrophoretic mobility shift assay (EMSA) was performed. The same 50 bp dsDNA fragment was used as in the nuclease assays. dsDNA (1 µM) was incubated at 22°C for 60 min with increasing concentrations of ComEA_Ng_ (0-200 µM) in 50 mM HEPES-NaOH pH 7.2, 50 mM NaCl, 5 mM MnCl_2_, 5% glycerol. Samples were separated by gel electrophoresis on 1% (w/v) agarose gels. The DNA was visualised using fluorescence illumination in a ChemiDoc imaging system (Bio-Rad).

#### Bioinformatic analysis

ComEC homologs were identified using BlastP (Blast v2.17.0) (Altschul *et al*, 1990) with the ComEC_Nc_ sequence as the query against the RefSeq Select database. The hits were ranked by query coverage, and the top 1000 Gram-positive and Gram-negative sequences were retained. Sequences were aligned using Clustal Omega (v1.2.4) (Sievers *et al*, 2011; Madeira *et al*, 2024), and alignment columns corresponding to gaps in the query sequence were manually removed. Phylogenetic analysis was performed with IQ-TREE 3 (Wong *et al*, 2025) to construct a maximum likelihood tree and estimate site-specific evolutionary rates. These rates were subsequently converted to the standard scale used by ConSurf (Ashkenazy *et al*, 2016) and mapped onto both the ComEC_Nc_ structure and the predicted ComEC-DNA translocation complex (ipTM: 0.66, pTM: 0.89).

The channel architecture of *apo* ComEC_Nc_ and the AlphaFold3-predicted translocation complex was analysed using MOLEonline (Raček *et al*, 2025; Abramson *et al*, 2024). Start and end points of the channel at the periplasmic entrance and the cytoplasmic exit regions were manually defined and a pore volume was generated. The predicted channel-lining residues were coloured according to conservation and mapped onto the structural models.

## Data availability

All data needed to evaluate the conclusions in this study are present in the paper and/or the Supplementary Information. The EM maps reported in this paper have been deposited in the Electron Microscopy Data Bank (EMDB) under accession codes EMD-57549 (ComEC competence and OB fold domains) and EMD-57548 (β-lactamase-like domain). Model coordinates have been deposited in the Protein Data Bank (PDB) under accession codes 30BU (ComEC competence and OB fold domains) and 30BT (β-lactamase-like domain).

## Author contributions

SD cloned constructs, purified proteins, performed nuclease activity and binding assays, prepared cryo-EM grids, assisted with cryo-EM data collection and processing and evaluated all data. DW processed cryo-EM data and built the model and evaluated data. TC and PA collected and processed cryo-EM data and TC built the model. MKH designed and supervised the study and wrote the manuscript with help from all authors. All authors contributed to figures.

## Acknowledgements

This work was supported by the Swiss State Secretariat for Education, Research and Innovation (SERI) under contract number MB22.00043 (M.K.H.), as well as the Swiss National Science Foundation (SNSF) grant 3200-0-239918 (M.K.H.). We are grateful to J. Rabl and A. Alexander for helpful discussions.

