## Supplementary Information for "Structural and biochemical insights reveal substrate-modulated nuclease activity of ComEC during DNA processing"

[3] ORCID: 0009-0000-4290-7792

[4] ORCID: 0000-0002-4337-4593

[5] ORCID: 0009-0004-4754-5830

[6] ORCID: 0000-0002-6353-6895

[7] ORCID: 0000-0003-1175-6826

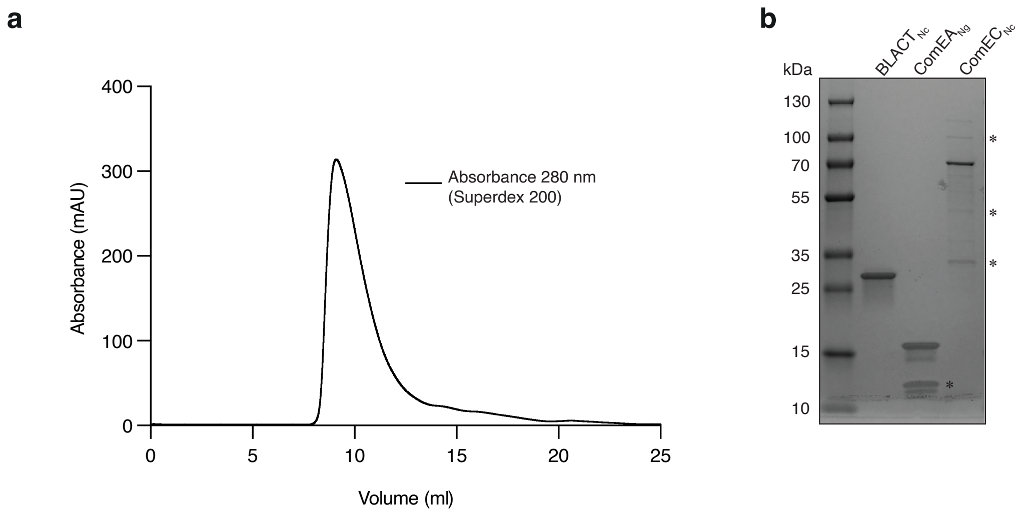


**Fig. S1: Purified proteins used in this study**

**a** Size exclusion chromatogram of LMNG-solubilised $ComEC_{Nc}$. **b** Purified proteins resolved by SDS-PAGE. ComEC and BLACT are derived from *N. carbonis* while ComEA is derived from *N. glycerini*. Additional bands (asterisks) observed in the ComEC and ComEA preparations were confirmed by mass spectrometry to correspond to degradation products and uncleaved sfGFP-ComEC. Further information about the constructs can be found in **Table S2**.

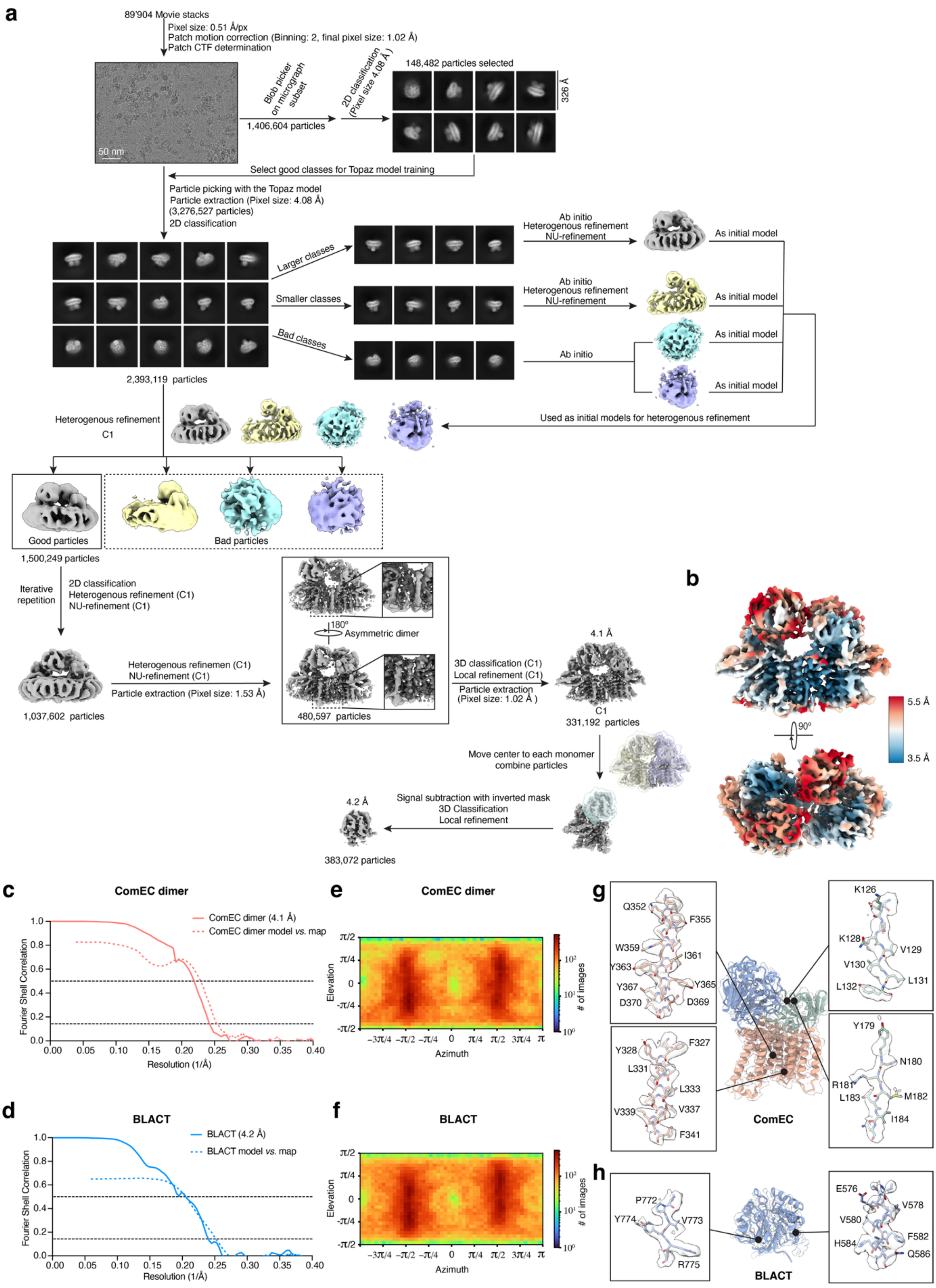

a
89'904 Movie stacks
Pixel size: 0.51 Å/px
Patch motion correction (Binning: 2, final pixel size: 1.02 Å)
Patch CTF determination
50 nm
Blob picker on micrograph subset
1,406,604 particles
2D classification (Pixel size 4.08 Å)
148,482 particles selected
326 Å
Select good classes for Topaz model training
Particle picking with the Topaz model
Particle extraction (Pixel size: 4.08 Å)
(3,276,527 particles)
2D classification
Larger classes
Smaller classes
Bad classes
Ab initio
Heterogenous refinement
NU-refinement
As initial model
2,393,119 particles
Heterogenous refinement
C1
Used as initial models for heterogenous refinement
Good particles
Bad particles
1,500,249 particles
Iterative repetition
2D classification
Heterogenous refinement (C1)
NU-refinement (C1)
180º
Asymmetric dimer
Heterogenous refinemen (C1)
NU-refinement (C1)
Particle extraction (Pixel size: 1.53 Å)
1,037,602 particles
480,597 particles
3D classification (C1)
Local refinement (C1)
Particle extraction
(Pixel size: 1.02 Å )
4.1 Å
C1
331,192 particles
Move center to each monomer
combine particles
Signal subtraction with inverted mask
3D Classification
Local refinement
4.2 Å
383,072 particles
b
5.5 Å
3.5 Å
90º
c
ComEC dimer
ComEC dimer (4.1 Å)
ComEC dimer model vs. map
Fourier Shell Correlation
Resolution (1/Å)
d
BLACT
BLACT (4.2 Å)
BLACT model vs. map
Fourier Shell Correlation
Resolution (1/Å)
e
ComEC dimer
f
BLACT
Elevation
Azimuth
### of images
g
Q352
F355
W359
I361
Y363
Y365
Y367
D370
D369
Y328
F327
L331
L333
V339
V337
F341
K126
K128
V129
V130
L131
L132
Y179
N180
R181
L183
M182
I184
ComEC
h
P772
Y774
V773
R775
BLACT
E576
V578
V580
H584
F582
Q586

**Fig. S2: Cryo-EM data processing and map validation for ComEC**

**a** Workflow of the cryo-EM data processing pipeline in CryoSPARC (Punjani *et al*, 2017). **b** Local resolution map showing that the competence domain is better resolved than the non-membrane domains. The weakest density was observed in the loop region connecting the competence domain and the β-lactamase-like domain. **c-d** Fourier shell correlation (FSC) (solid) and model vs. map FSC (dashed) curves for the cryo-EM reconstructions of the ComEC dimer (**c**) and the β-lactamase-like domain (BLACT) (**d**). **e-f** Euler angle distributions of particles used for the final reconstructions of the ComEC dimer (**e**) and the β-lactamase-like domain (**f**). **g-h** Representative regions of density in the ComEC dimer (**g**) and β-lactamase-like domain (**h**) reconstructions, illustrating map quality and fitted atomic models.

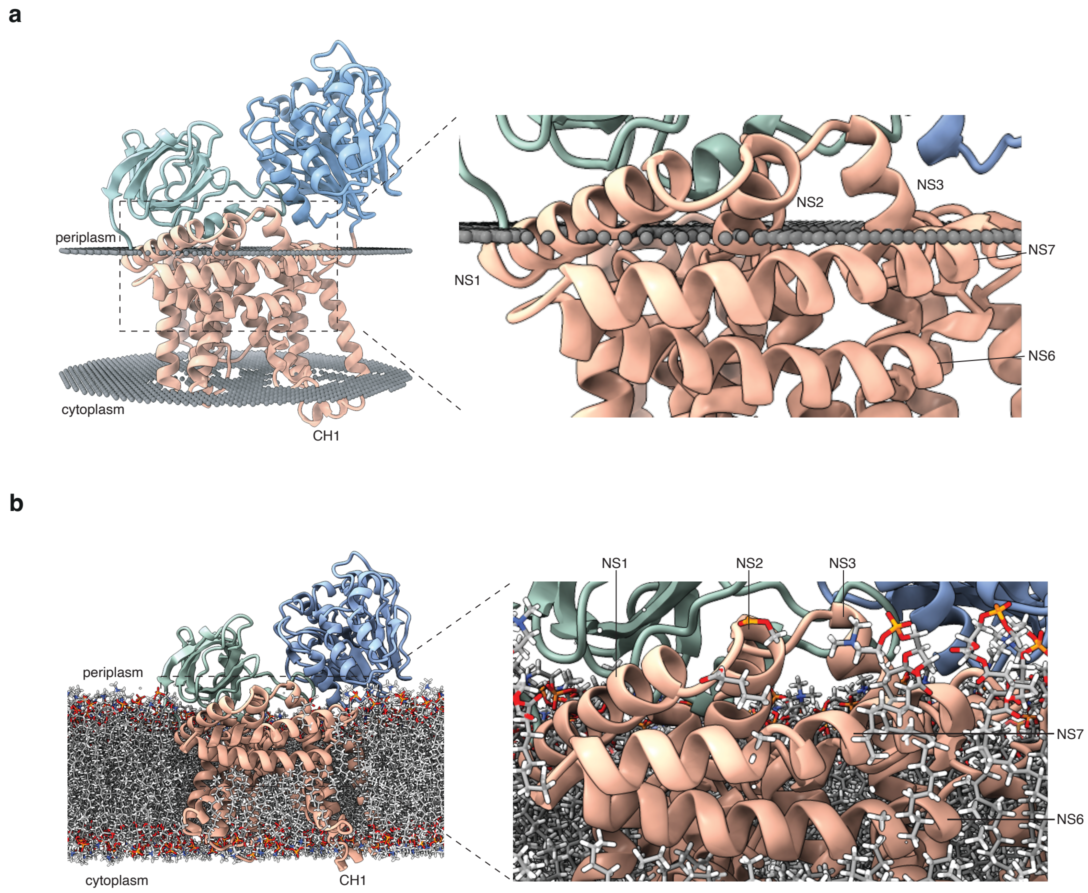


**Fig. S3: Orientation of ComEC in a lipid bilayer**

**a-b** Predicted insertion of ComEC into a lipid bilayer using PPM 3.0 (Lomize *et al*, 2022) (**a**) and CHARMM-GUI v3.7 (Jo *et al*, 2007, 2009) (**b**). Close-up views show the positioning of NS1 (amphipathic), NS2–3 (periplasmic), and NS6–7 (membrane-embedded) relative to the lipid bilayer.

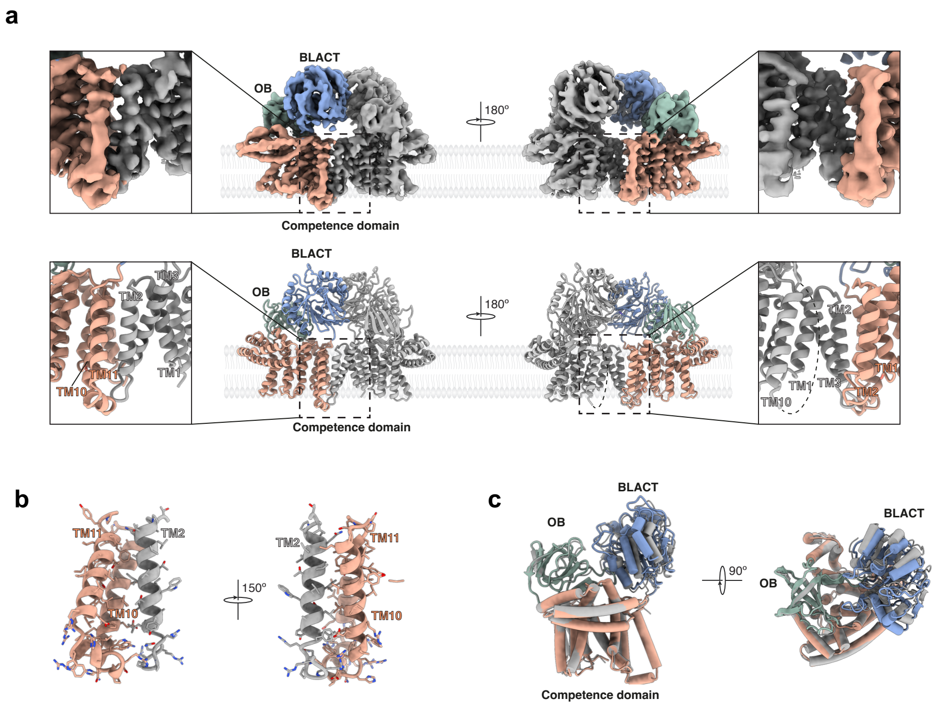


**Fig. S4: Cryo-EM reveals an asymmetric ComEC dimer in detergent micelles**

**a** Cryo-EM map (top) and corresponding atomic model (bottom) of the ComEC dimer, showing that dimerisation is mediated by an asymmetric interaction between three transmembrane helices within the competence domains. **b** Close-up views of the dimer interface highlighting hydrophobic interactions between TM10 and TM11 of one protomer and TM2 of the opposing protomer. **c** Superposition of the two ComEC protomers within the dimer, showing that the β-lactamase-like domain exhibits greater flexibility compared to the competence domain and OB fold. Two views are shown for each panel.

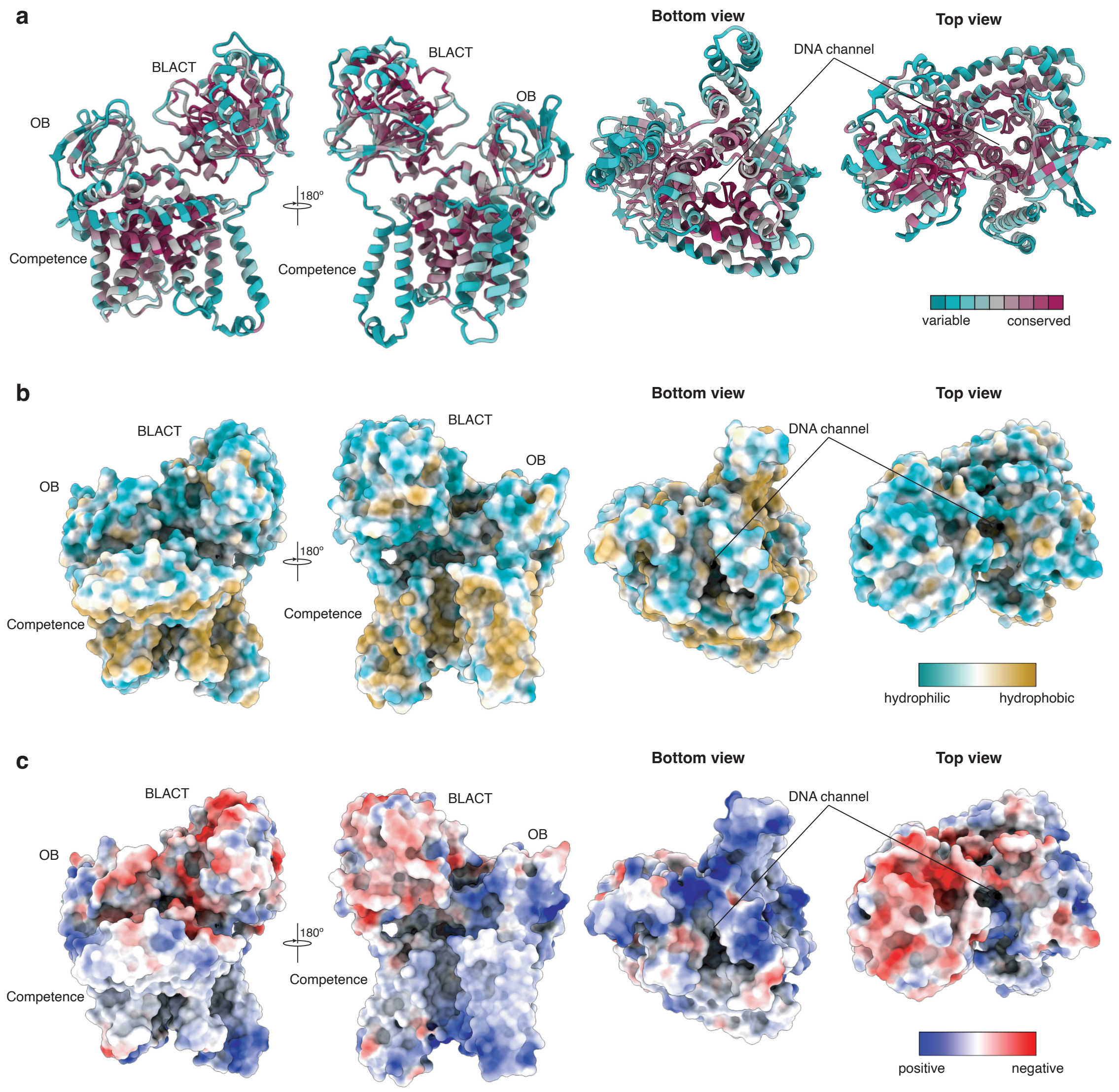


**Fig. S5: Conservation, hydrophobicity and electrostatic potential of ComEC$_{Nc}$**

**a-c** ComEC$_{Nc}$ coloured according to sequence conservation (cyan, low; purple, high) (**a**), hydrophobicity (cyan, hydrophilic; yellow, hydrophobic) (**b**), and electrostatic potential (blue, positive; red, negative) (**c**). The three domains and the DNA channel are indicated. For each panel, two side views as well as top and bottom views are shown.

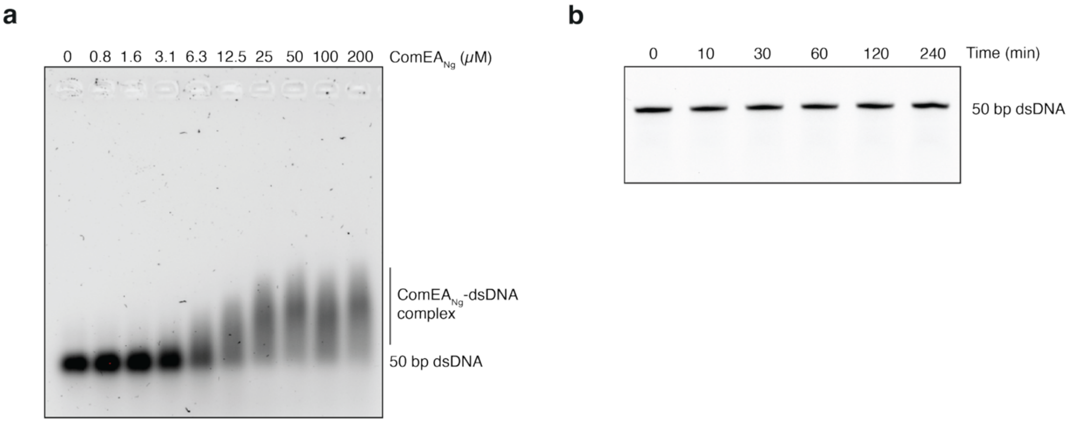


**Fig. S6: ComEA$_{Ng}$ binds, but does not degrade dsDNA *in vitro***

**a** Electrophoretic mobility shift assay (EMSA) showing binding of ComEA$_{Ng}$ to dsDNA. A 50 bp dsDNA fragment (1 μM) was incubated with increasing concentrations of purified ComEA$_{Ng}$ (0-200 μM) comprising the oligomerisation and DNA-binding domain. DNA was labelled at the 5' end with a FAM fluorophore. **b** Time-course nuclease assay showing that ComEA$_{Ng}$ alone does not exhibit detectable endonucleolytic activity under the conditions used in **Fig. 4c**.

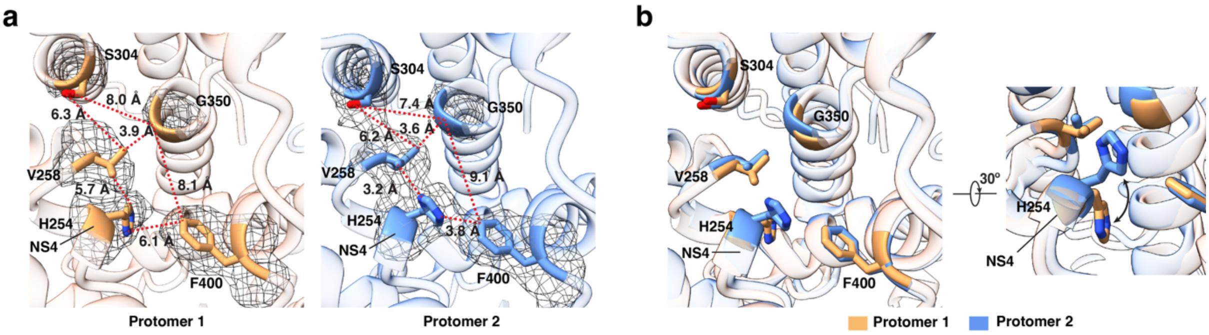


**Fig. S7: Comparison of the periplasmic channel entrance between ComEC protomers in the dimer**

**a** Close-up view of the region around NS4 in both protomers of the ComEC dimer structure, showing key side chains with corresponding EM density. Distances between selected side chains are indicated. **b** Superposition of the two protomers highlighting conformational differences in the H254 side chain.

#### Table S1: Data collection, 3D reconstruction and model statistics

| | ComEC dimer | BLACT |
|---|---|---|
| **Data collection** | | |
| Micrographs | 89,904 | |
| Voltage (kV) | 300 | |
| Detector | K3 camera | |
| Pixel size (Å/pixel) | 0.506 | |
| Defocus range (μm) | -1.1 to -2.7 | |
| Electron exposure (e-/Å²) | 53 | |
| Software | EPU (v. 3.10.0) | |
| **Reconstruction** | | |
| Software | CryoSPARC (v. 4.7.1) | |
| Final particles (No.) | 331,192 | 383,072 |
| FSC threshold | 0.143 | |
| Resolution (Å) | 4.13 | 4.17 |
| Symmetry | C1 | |
| Map sharpening B-factor (Å²) | -180 | -150 |
| **Model building** | | |
| Software | Coot (v. 9.8.5) | |
| Model composition | | |
| Residues | 1024 | 273 |
| Non-hydrogen atoms | 7929 | 2097 |
| **Refinement** | | |
| Software | Phenix (V. 2.0) and RosettaCM | |
| R.M.S deviations | | |
| Bond length (Å) | 0.002 | 0.002 |
| Bond angles (°) | 0.562 | 0.444 |
| Validation | | |
| MolProbity score | 1.38 | 1.42 |
| Clashscore | 6.91 | 7.67 |
| Rotamer outliers (%) | 0.00 | 0.00 |
| Ramachandran plot | | |
| Favored (%) | 98.82 | 98.89 |
| Allowed (%) | 1.18 | 1.11 |
| Outliers (%) | 0.00 | 0.00 |
| Masked CC | 0.71 | 0.65 |

**Table S2: Plasmids used in this study**

| Name | Relevant genotype/description | Source/Reference |
|---|---|---|
| pBAD | *E. coli* expression vector, pBAD promoter, AmpR | |
| pBAD-sfGFP-3C-StrepII | *E. coli* expression vector, N-terminal sfGFP tag, C-terminal StrepII tag, pBAD promoter, AmpR | |
| pBAD-sfGFP-3C-ComEC$_{Nc}$-StrepII | *N. carbonis* wild-type *comEC* | This study |
| pOPINS | *E. coli* expression vector, N-terminal His$_6$-SUMO tag, T7 promoter, KanR | (Assenberg *et al*, 2008) |
| pOPINS-BLACT$_{Nc}$ | *N. carbonis* β-lactamase-like domain (532-801) | This study |
| pET28a(+) His$_6$-SUMO ComEA$_{Ng}$ | *E. coli* expression vector, N-terminal His$_6$-SUMO tag, T7 promoter, KanR, *N.glycerini comEA* (oligomerisation and DNA-binding domain) | This study |

#### Supplementary Table 3: Oligonucleotides used in this study

| Name | Sequence (5' to 3') | Construct |
|---|---|---|
| *Cloning* | | |
| pOPINS-linearise-BLACT$_{Nc}$-Fw | CTATTCCGGGCGGTTACTGATAA AGCTTTCTAGACCATTTAAACAC CACCACC | pOPINS fragment for insertion of BLACT$_{Nc}$ |
| pOPINS-linearise-Rv | ACCACCGATCTGTTCGCGATGC | pOPINS fragment |
| pOPINS-BLACT$_{Nc}$-Fw | CGAACAGATCGGTGGTCAAGAA GAGCTGAAAGTCACATTCATAGA TGTGGG | pOPINS-BLACT$_{Nc}$ |
| pOPINS-BLACT$_{Nc}$-Rv | GTAACCGCCCGGAATAGTTGTGT CCAC | pOPINS-BLACT$_{Nc}$ |
| pBAD sfGFP-3C-StrepII-linearise-Fw | GGTTACTGGAGCCACCCGCAGT TCGAAAAGTGAAGTGGATGGAG CCACCCG | pBAD fragment for insertion of ComEC$_{Nc}$ |
| pBAD sfGFP-3C-StrepII-linearise-Rv | ACCACCAGAAGAGCCGCCTGGA CCTTGAAACAAAACTTCTAAACC AGATTTGTAGAG | pBAD fragment for insertion of ComEC$_{Nc}$ |
| pBAD-ComEC$_{Nc}$-Fw | GCGGCTCTTCTGGTGGTATAGC GGCCCCGCTGGTG | pBAD-ComEC$_{Nc}$ |
| pBAD-ComEC$_{Nc}$-Rv | GGTGGCTCCAGTAACCGCCCGG AATAGTTGTGTCC | pBAD-ComEC$_{Nc}$ |
| *Biophysical assays*[#] | | |
| 5'Phos-5'FAM*-Fw | [p]T*GGGTAGGATCATCAGTAATAA GGATAGTGGGAAAGCTCACAGA CCACCT | Nuclease assay |
| 5'Phos-5'FAM*-(PTO) Fw | [p]T*/G/G/G/T/A/G/G/A/T/C/A/T/C/A/ G/T/A/A/T/A/A/G/G/A/T/A/G/T/G/G/ G/A/A/A/G/C/T/C/A/C/A/G/A/C/C/A/ C/C/G | Nuclease assay |

[#] dsDNA substrates were generated by annealing the labelled strand to a complementary 5′-phosphorylated, unlabeled strand. All DNA probes were 50 nt in length.

* FAM was attached to the first base (thymine, T) at the 5' terminus.

[p] 5' phosphate modification.

/ Cleavable phosphodiester bond.

/ Uncleavable phosphorothioate bonds.
